# Adhesion and injury cues enhance blackworm capture by freshwater planaria

**DOI:** 10.1101/2025.04.09.647989

**Authors:** Ishant Tiwari, Hiteshri Chudasama, Harry Tuazon, Saad Bhamla

## Abstract

In aquatic ecosystems, freshwater planarians (*Dugesia spp*.) function as predators, employing specialized adaptations for capturing live prey. This exploratory study examines the predatory interactions between the freshwater planarian *Dugesia spp*. and the California blackworm (*Lumbriculus variegatus*). Observations demonstrate that *Dugesia* is capable of capturing prey more than twice its own length. The predation process involves a dual adhesion mechanism whereby the planarian adheres simultaneously to the blackworm and the substrate, effectively immobilizing its prey. Despite the rapid escape response of blackworms, characterized by a reversing spiral swimming gait, planarian adhesion frequently prevents successful escape, although notably larger blackworms exhibit increased escape success. Subsequently, *Dugesia* employs an eversible pharynx to initiate ingestion, consuming the internal tissues of the blackworm through suction. Injury in blackworms emerged as a significant predictor of predation events, suggesting the potential involvement of chemical cues in prey detection, although this warrants further investigation. This study provides insights into the biomechanics and behaviors of predation involving two interacting muscular hydrostats, highlighting the critical adaptations that enable planarians to subdue and consume relatively large, mobile prey.

## Introduction

Brown freshwater planarians, particularly *Dugesia spp*., exhibit diverse predatory behaviors, feeding on a variety of prey including aquatic oligochaetes, insect larvae, and crustaceans (Perich and Boobar 1990; Vila-Farré and C Rink 2018). Serving as predators in freshwater habitats such as rivers, streams, and ponds, they provide an opportunity to explore ecological themes like interspecies competition and predation strategies (Sluys and Kawakatsu 2005). Morphologically, *Dugesia spp*. are characterized by their small body size (typically under two centimeters), distinctive brown coloration, flattened dorsoventral body shape, and twin ocular spots (Fig.1A). They possess regenerative capabilities, enabled by abundant stem cells known as neoblasts, allowing complete regeneration of lost tissues and organs Baguñà et al. (1990).

**Fig. 1.**
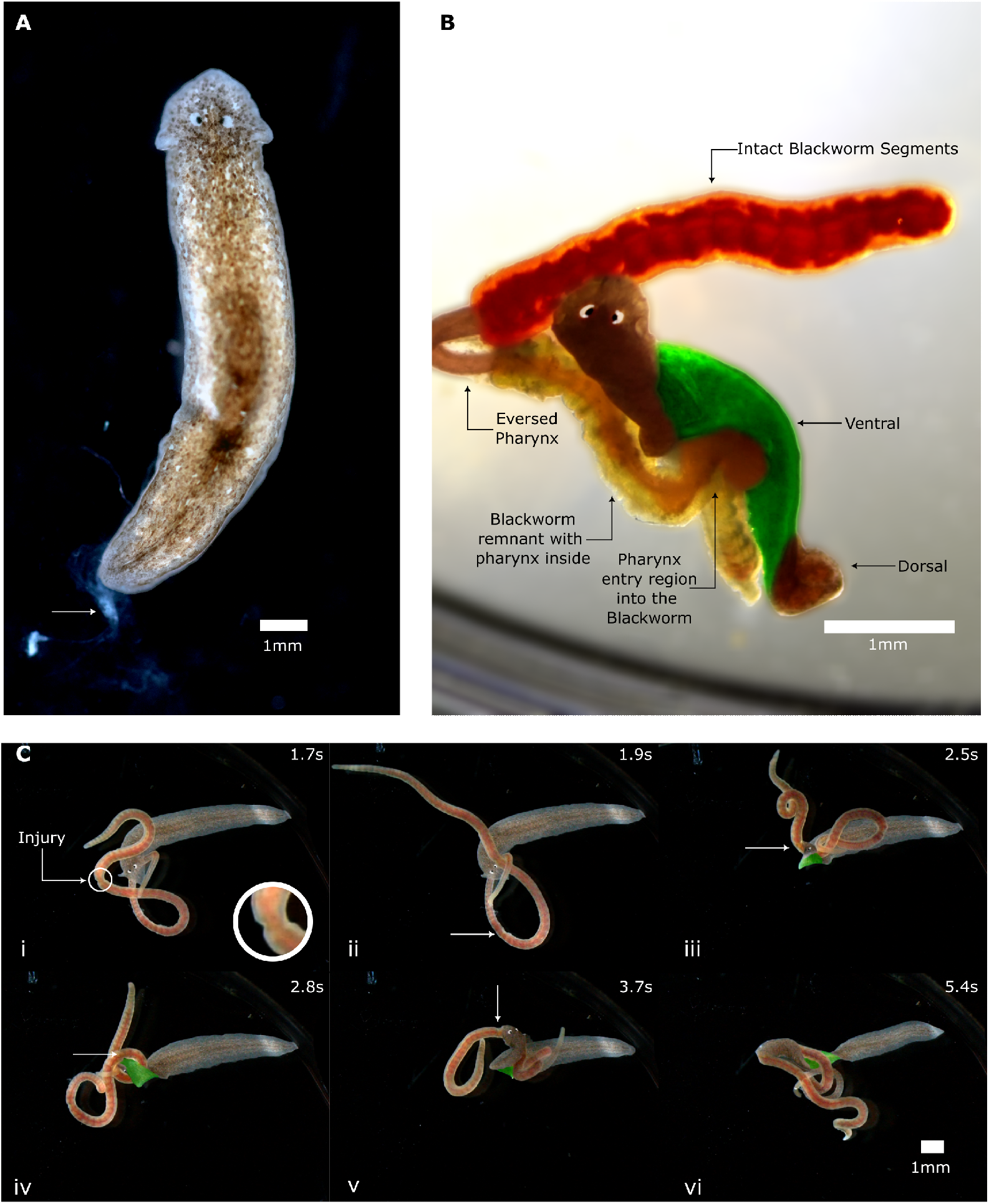
Planarian morphology, feeding, and injury-targeted adhesion. (A) Dorsal view of *Dugesia tigrina* showing auricles, paired eyespots, and adhesive mucus visible at the posterior end (white arrow). (B) False colored snapshot of *Dugesia dorotocephala* feeding on a blackworm. The planarian body is twisted, exposing both dorsal (brown) and ventral (green) surfaces. The everted pharynx (brownish-red) leaves behind a collapsed, emptied segment of the blackworm (yellow) surrounding the pharynx, with the pharynx now progressing toward an intact region (red). Supplementary Video 1 contains the full interaction without false coloring. (C) Time series (i–vi) from high-speed video (1000 FPS, Supplementary Video 2) showing *D. tigrina* interacting with an injured blackworm. The injury site is marked (panel i shows zoomed inset of the injury), and the planarian is seen contacting and twisting around the worm, revealing ventral regions as it anchors near the injury site. This sequence illustrates adhesive contact and behavioral responses to localized damage.

The predatory behavior of freshwater planarians involves specialized strategies adapted to capturing agile prey. For example, they demonstrate predation on mosquito larvae through direct physical contact and mucus adhesion (Perich and Boobar 1990). When targeting California blackworms (*Lumbriculus variegatus*), planarians face substantial biomechanical challenges due to the blackworms’ robust escape response, characterized by a rapid reversing spiral swimming gait (Drewes and Fourtner 1989; Drewes 1990). Additionally, blackworms employ defensive collective behaviors, forming entangled aggregates or “worm blobs,” significantly larger than individual predators (Patil et al. 2023; Tuazon et al. 2022, 2023).

Despite the blackworm’s well-documented escape responses including rapid helical locomotion and the formation of entangled collectives, interactions between *Dugesia* and *Lumbriculus variegatus* and the corresponding biomechanics are still underexplored. These two organisms often inhabit the same freshwater environments, including shallow streams, ponds, and leaf-littered substrates, where physical proximity may increase the likelihood of predator-prey encounters (Vila-Farré and C Rink 2018; Pennak 1978; Boddington and Mettrick 1974; Pickavance 1971). The shared ecological context provides a basis for exploring how prey defenses influence predator behavior in such benthic microhabitats.

Here, we study the behavioral and physical interactions between *Dugesia* and *L. variegatus*, focusing on the capture phase of predation. Using observational data, we characterize how the planarian initiates and sustains contact with the blackworm and examine how this engagement affects the worm’s ability to initiate escape responses. We also investigate whether injury may play a role in triggering predation. These observations are interpreted in the context of muscular hydrostat function, a category that includes both organisms due to their reliance on deformable, muscle-based structures without rigid skeletons (Kier and Smith 1985; Miyamoto et al. 2020). The interaction provides a tractable model for studying how flexible-bodied invertebrates engage during antagonistic contact in aquatic settings.

## Materials and methods

### Animals

We sourced the brown planaria (*Dugesia tigrina* and *Dugesia dorotocephala*) from two sources; Flinn Scientific (*Dugesia tigrina*) and Home Science Tools (*Dugesia dorotocephala*). We obtained the California blackworms (*Lumbriculus variegatus*) from Ward’s Science. Both organisms were reared in a plastic box (35 × 20 × 12 cm) with filtered water at room temperature (∼21^°^C). Gentle aeration was provided using aquarium air stones to maintain oxygen levels. The water was exchanged daily to ensure its quality, and the worms were fed crushed tropical fish pellets twice a week. Planaria (*Dugesia dorotocephala* and *Dugesia tigrina*) were housed in covered containers under the same water maintenance and aeration protocol. They were starved for seven days prior to the experiments to standardize hunger levels. Since our study involved working with only invertebrates, it did not require an Institutional Animal Care and Use Committee (IACUC) approval.

### Imaging

Behavioral interactions between planaria and blackworms were recorded using an ImageSource DFK 33UX264 camera (Charlotte, NC) positioned above the arena, at 30 FPS.

For high-speed recordings, we used a Leica MZ APO stereomicroscope (Heerbrugg, Switzerland) coupled with a Chronos 2.1 high-speed camera (Kron Technologies, Canada), capturing at 1000 frames per second (FPS). A gooseneck light source was used to illuminate the arena during both sets of recordings. We made a total of N=6 high speed (1000 FPS) recordings of the planaria blackworm interaction, using distinct blackworms and planaria.

### Behavioral Testing

The arena consisted of a 35 mm petri dish containing filtered water prepared using a reverse osmosis (RO) system. Salts (NilocG Aquatics REKHARB carbonate hardness (KH) booster and NilocG Aquatics REGEN general hardness (GH) booster) were added to achieve spring water conditions with a total dissolved solids (TDS) concentration of approximately 50 ppm, following the manufacturer’s guidelines. Four behavioral experiments were conducted: (1) injured long worms with planaria, (2) injured short worms with planaria, and two control conditions with uninjured worms (long and short). Blackworms were injured by gently pinning them between a plastic pipette and the substrate, resulting in a noticeable kink in their bodies. Healthy blackworms (N=10 long worms: 30.1 ± 3.3 mm, N=10 short worms: 12.1 ± 3.7 mm contour length) and planaria (N=20, 12.6 ± 2.9 mm contour length) were randomly selected for each trial. Planaria were placed in the petri dish with 5 mm depth of prepared water for 30 minutes prior to introducing the blackworm. Each interaction was recorded immediately upon contact between planarian and blackworm, and videos lasted approximately 30 minutes. Each experiment was repeated for N = 5 replicates. This meant a total of 20 behavioral trials; 4 Groups *×* N=5 independent replicates.

The “grip and wrap” behavior was defined as any action in which the planarian adhered to the blackworm using its body and then coiled around it. The gripping behavior was further described into sub-events: substrate grip and prey grip. In substrate grip, the planarian anchored its posterior end onto the petri-dish substrate, while its anterior end reached towards the blackworm. In prey grip, the planarian directly adhered to the worm’s body, maintaining contact as the worm attempted escape. Following successful gripping, the planarian wrapped itself around the tube-like geometry of the blackworm as the planarian everted its pharynx for feeding. Predation was classified as successful if the planarian successfully fed on the worm. All behaviors were documented and categorized using video analysis according to these criteria.

### Data Analysis

We tracked body positions from high-speed recordings using DLTdv8 Hedrick (2008), a MATLAB-based digitizing tool designed for kinematic analysis. In this study, we used DLTdv8 for manual frame-by-frame tracking. For blackworms, we marked the head and tail tips, which served as clean and reliable landmarks. For planaria, we tracked one eyespot and the posterior tail tip. Tracking was skipped, When points were occluded due to body overlap or contact during behavior.

To obtain contour length measurements, we imported still frames from each trial into ImageJ and traced the body outlines. These measurements were used to estimate individual length scales and categorize blackworms as either “short” (< 20 mm) or “long” (≥ 20mm) for experimental grouping. Tracking and length data were used to analyze body positioning and relative size ratios (worm length to planarian length) during predator-prey interactions.

To create the composite shown in Figure 2A, we used a frame-wise pixel-wise maximum intensity projection over the 2–5 second interval of the high-speed video. Each pixel in the final image corresponds to the brightest intensity recorded at that spatial location during this window. To indicate temporal progression, we assigned a hue to each frame based on its timestamp, shifting from red to blue across the sequence.

**Fig. 2.**
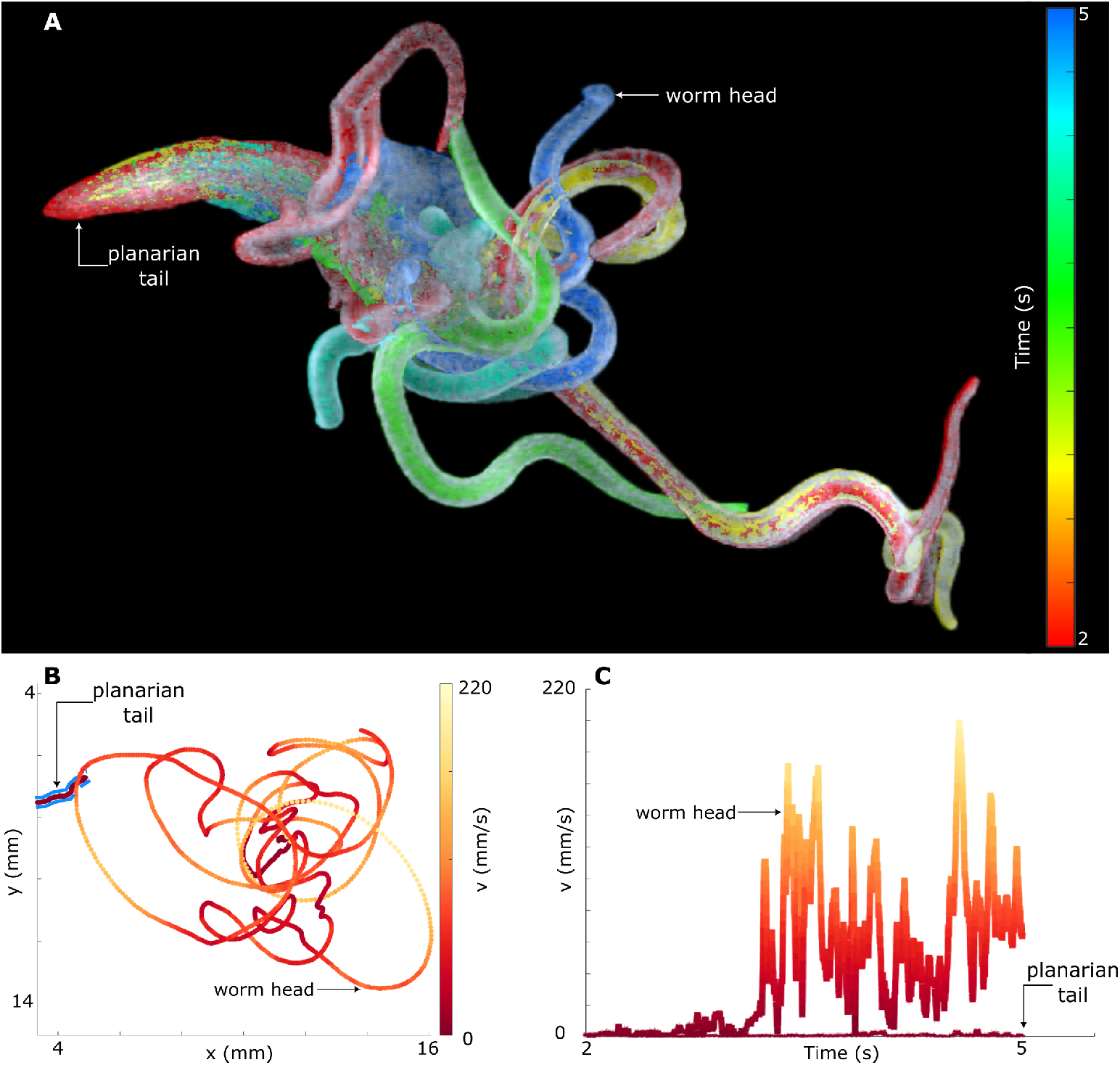
Planarian-substrate adhesion during blackworm escape attempts. (A) Overlay of high-speed video frames showing a blackworm’s rapid spiral escape response during a predation event with *Dugesia tigrina*. Frame hue transitions from red (early) to blue (later), representing time progression over a 3-second interval (2–5 seconds). Multiple blackworm body postures indicate high mobility, while the planarian tail remains localized in the upper left, suggesting strong adhesion to the substrate. (B) Trajectories of the blackworm head and planarian tail from panel A. Each point is color-coded by instantaneous speed, with yellow indicating high (∼ 200mm/s) and red indicating low speed (Median = 0.74mm/s). The planarian’s trajectory (outlined in blue) is compact (4 < *x* < 6 mm, 6 < *y* < 8mm) and slow, while the blackworm shows high-speed (Median = 31.77mm/s), large-displacement motion (4 < *x* < 16 mm, 4 < *y* < 14mm). (C) Instantaneous speeds of the worm head and planarian tail plotted together. The blackworm reaches speeds up to 200 mm/s, while the planarian tail speed remains near zero. These panels highlight the planarian’s use of posterior adhesion to resist escape forces from an actively struggling prey. See Supplementary Video 3 for the full interaction.

### Statistical Analysis

To assess whether worm length affected escape outcomes, we compared the length ratios (worm length relative to planarian length) of trials that resulted in successful escape versus those that did not. We performed a Mann–Whitney U test (also known as the Wilcoxon rank-sum test) to test for differences between the two groups Mann and Whitney (1947). As a complementary nonparametric effect size measure, we computed Cliff’s Delta along with 95% confidence intervals using bootstrapping (10,000 iterations) Cliff (1993). All analyses were conducted in MATLAB.

## Results

### Planarian morphology and adhesive behavior

Figure 1A shows a dorsal view of *Dugesia tigrina*, with visible auricles and paired eyespots near the anterior end Hyman (1939). A white substance is present at the posterior end (white arrow, Figure 1A), corresponding to mucus secretions deposited during movement. These secretions are associated with adhesion to substrates and prey Pickavance (1971); Perich 165 and Boobar (1990), and appear to assist the planarian in 166 maintaining a stable grip during behavioral interactions.

### Feeding via everted pharynx

Figure 1B presents a snapshot of *Dugesia dorotocephala* feeding on a captured blackworm. The planarian’s body is twisted along its long axis as it adheres to different regions of the blackworm, revealing both dorsal (brown) and ventral (green) surfaces in this top-down view. The everted pharynx (brownish red) is inserted into a collapsed segment of the blackworm (yellow), which has been depleted of internal contents. A more distal segment of the blackworm remains intact (red). The pharynx has already passed through the emptied region and is entering the next, leaving behind a dead husk that surrounds the pharynx externally. The complete sequence of this interaction is included in **SI video 1**. Previous work describes the planarian pharynx as a muscular hydrostat capable of coordinated movement through the pharyngeal nervous system Miyamoto et al. (2020). The observed feeding posture illustrates progression of pharyngeal activity along the worm’s internal cavity, leaving behind a collapsed cuticle that envelops the feeding structure.

### Adhesive engagement and response to injury

Figure 1C shows a time series (i–vi) of *Dugesia tigrina* interacting with an injured blackworm. These high-speed snapshots, extracted from a recording provided in **(SI Video 2)**, follow a planarian as it explores the surface of a blackworm displaying a visible injury site. In panels i and ii, the planarian contacts the tail region of the worm, while the injury point remains separate. As the worm initiates escape movements, the planarian shifts position and approaches the injury site. In panels iii through vi, the planarian attaches near this region while undergoing a 360-degree twist along its longitudinal axis. This twisting motion, likely driven by simultaneous escape behavior in the blackworm and active adhesion by the planarian, reveals ventral surfaces in a dorsal view. These ventral regions are false-colored green in the images for easier visualization.

The sequence in Figure 1C visually demonstrates the adhesive capacity of the planarian body and its ability to maintain contact with moving prey. The targeted attachment near the injury site suggests a possible behavioral sensitivity to damaged tissue. Similar targeting behaviors have been observed in other soft-bodied predators Kutschera (2003), and injury-directed attraction in planarians has been hypothesized Pickavance (1971).

### Adhesive anchoring during prey capture

Figure 2A presents an overlay of high-speed (recorded at 1000 FPS) video frames capturing a predation event involving *Dugesia tigrina* and a blackworm. Frame colors progress from red (earlier) to blue (later), representing a three-second interval (2 to 5 seconds). The blackworm exhibits pronounced helical twisting movements as it attempts escape, reflected in the multiple body postures visible across the frame. In contrast, the tail of the planarian remains confined to a small region in the upper left corner, suggesting stable adhesion to the substrate despite significant prey movements.

In Figure 2B, trajectories of the planarian tail and blackworm head positions from panel A are shown. Points along each trajectory are color-coded according to instantaneous speed, with higher speeds (∼ 200mm/s) in yellow and lower speeds (∼ 0mm/s) in red. The planarian’s tail trajectory (outlined in blue) occupies a compact region (4 < *x* < 6,6 < *y* < 8mm) with consistently low speeds (∼ 0mm/s), while the worm’s head trajectory spans nearly the entire frame (4 < *x* < 16,4 < *y* < 14mm), demonstrating extensive motion due to escape behaviors.

Figure 2C compares instantaneous speeds of the blackworm head and planarian tail. While the blackworm achieves speeds approaching 200 mm/s, the planarian’s tail speed remains negligible (∼ 0 mm/s). Together, these three panels demonstrate that *Dugesia* effectively utilizes adhesion to anchor itself firmly to the substrate, counteracting the prey’s vigorous escape attempts. The complete video sequence of this event is available in **(SI Video 3)**.

### Injury and size influence planarian predation outcomes

In panel 3A, a planarian and blackworm interaction is depicted with contour lengths marked by dotted lines to define worm and planarian lengths. An inset highlights a kinked section on the blackworm body labeled “Injury?”, indicating the experimental condition of injury tested in specific trials. The figure visually demonstrates two measurable events: “Attack time,” defined as the duration from initial approach to the planarian’s attempt at attachment, and “Escape?” indicating the worm’s success or failure in evading the planarian after attack.

Figure 3B presents a scatter plot of planarian length (y-axis) against worm length (x-axis), with each point colored red if an attack occurred and blue if there was no attack. The distribution indicates that trials covered a diverse range of predator-prey length combinations across 20 total trials (four experimental groups, *N* = 5 replicates each).

**Fig. 3.**
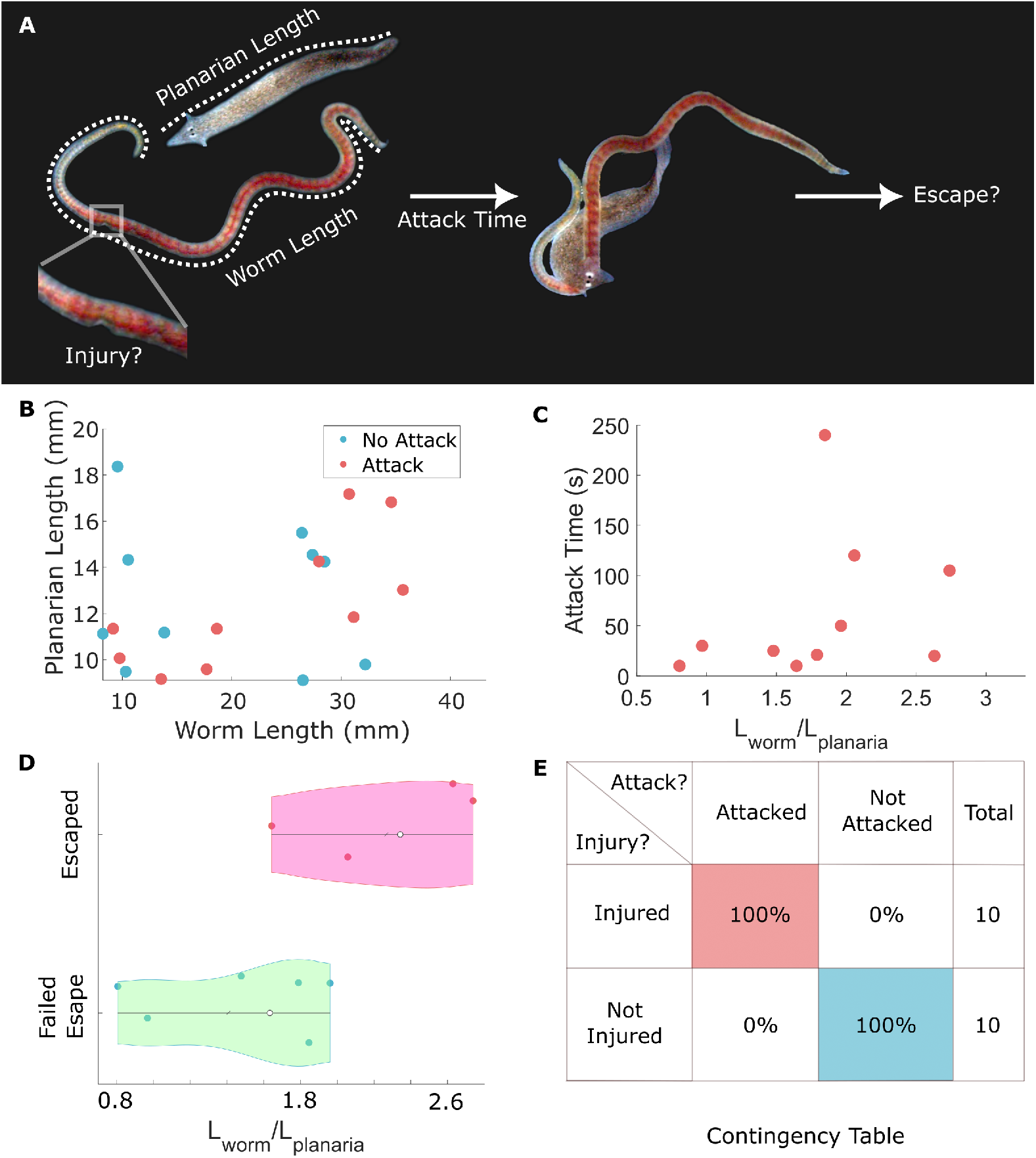
Behavioral assays reveal injury targeting and size-dependent escape outcomes. (A) Schematic overview of a behavioral trial. Dotted lines mark the contour lengths of a blackworm and a planarian. Zoomed inset identifies a body kink as an injury, indicating the test condition for certain trials. Arrows show the timeline of observed behaviors: “Attack time” marks the interval between planarian approach and attachment, followed by the boolean outcome (Success/Failure to escape, see panel D) labeled “Escape?”. (B) Scatter plot of planarian length versus worm length across all 20 trials. Each point is red if an attack occurred and blue otherwise. The spread indicates a range of relative predator-prey sizes across experimental conditions. (C) Attack time plotted against the length ratio (*L*_worm_*/L*_planaria_). Most attacks occur within 50 seconds (median = 27.5 s, *N* = 10). (D) Violin plots show the relationship between length ratio and escape outcome. Pink violins correspond to successful escapes, green to failed escapes. Worms with higher length ratios (median ratio *L*_worm_ /*L*_planaria_ =2.3) more often escape (Mann–Whitney U test: *p* = 0.0667; Cliff’s delta = 0.75 (95% CI: 0.167–1), suggesting a strong effect despite small sample size.) White dot in the center indicates median of the group. (E) Contingency table comparing injury status and attack outcome. All injured worms were attacked; none of the uninjured worms were, regardless of their lengths (N=20 total experiments).

Figure 3C plots “attack time” against the length ratio (worm length divided by planarian length). Attack times generally cluster below 50 seconds (median = 27.5 s, *N* = 10 attacks).

Figure 3D uses horizontal violin plots to investigate the relationship between length ratio (*L*_worm_*/L*_planaria_) and escape probability. Green represents worms that failed to escape, while pink indicates successful escapes. Visually, worms that escaped exhibit higher length ratios (median ratio *L*_worm_*/L*_planaria_ =2.3; *N* = 4 escapes out of 10 attacks). A Mann–Whitney U test comparing escaped and non-escaped groups yields a *p*-value of 0.0667, suggesting no statistically significant difference at traditional thresholds. However, considering the limited sample size, Cliff’s delta analysis indicates a strong effect (delta = 0.75; 95% CI: 0.167 to 1), suggesting a meaningful relationship where worms with greater length ratios tend to escape more successfully.

Figure 3E provides a contingency table examining the relationship between worm injury and planarian attack probability. Injury emerges as a clear predictor of attack occurrence, with 100% of injured worms attacked and no uninjured sworms attacked, irrespective of their lengths.

These results collectively suggest that the planarian’s predation behavior strongly exploits prey injury cues and that larger prey relative to the predator size show greater escape success, possibly highlighting physical limitations on prey handling.

## Discussion

Our study characterized predator-prey interactions between freshwater planarians and blackworms, highlighting planarians’ use of adhesive secretions and targeted attacks on injured worms. Using high-speed videography, we captured detailed behavioral sequences demonstrating how planarians immobilize prey by simultaneously adhering to the substrate and their target. We showed that injury significantly increased vulnerability, with all injured blackworms being targeted irrespective of their size, while uninjured worms were never attacked during trials.

Previous studies have documented similar interactions among freshwater planarians and oligochaete worms. Pickavance (1971) observed that *Dugesia tigrina* selectively preyed upon injured individuals of *Lumbriculus*, indicating an opportunistic feeding strategy in benthic environments. Additionally, previous research Boddington and Mettrick (1974) reported another *Dugesia* species preying frequently on tubificid worms, further supporting the ecological significance of these interactions. Our results build on these findings by visually demonstrating the biomechanics of these interactions, specifically planarians’ effective use of adhesive mucus to subdue active prey.

Despite providing clear behavioral evidence, our study has limitations. All experiments were conducted in simplified laboratory arenas with smooth glass or plastic substrates, which reduced the blackworm’s ability to burrow or hide, strategies commonly employed in natural sediments. In more structurally complex environments, such as leaf litter or fine-grained detritus, prey may evade or resist planarian attacks more effectively, potentially altering the observed outcomes. However, the use of clean, flat substrates allowed unobstructed visualization of subtle contact behaviors, body orientation changes, and adhesive responses that may be difficult to resolve in more ecologically realistic settings. Future experiments incorporating granular or heterogeneous substrates, as well as varying environmental parameters like conspecific density, could help contextualize these results and assess their generality.

The established experimental paradigm involving interaction3s74 between two muscular hydrostats, planarian and blackworm, creates several openings for future work. Moreover, the pharyngeal ability to evert, navigate along prey tissue, and draw in material suggests coordination across multiple mechanical and sensory subsystems Miyamoto et al. (2020). High-resolution 3D imaging of the planarian, combined with mechanical modeling of a ribbon like structure wrapping around a cylindrical worm is another research direction that lies at the interface of biomechanics and geometry. The adhesive system itself remains poorly understood. The effective adhesive properties in a hydrated environment raise questions about the underlying physical or biochemical mechanisms and whether these can be modulated in response to substrate/prey.

Behaviorally, questions remain about prey specialization. While this study focused on worm capture, comparisons with how planarians handle other prey types may reveal trade-offs or constraints of their flexible, elongate body plan. A comparative analysis could test whether the observed strategies are general-purpose or specifically suited to long, flexible prey.

Finally, investigating how planarians respond to collectives of blackworms would be an interesting problem involving the structural mechanics of the planarian’s ribbon shape and the topological mechanics of an entangled worm blob. Previous research involving another invertebrate predator, *Helobdella spp*. leech, reported a complex struggle between the spiral entombment strategy of the leech and the entangled blob formation of the blackworms Tuazon et al. (2024). In that work, the leech faced a challenge of pulling a single worm out of an entangled blob of worms. With the adhesive grip and wrap strategy of the planarian, whether adhesion and wrapping scale effectively to such entangled, multibody targets could extend the findings of this work into more complex and ecologically representative settings.

## Supplementary data

Supplementary data available at *ICB* online.

## Author contributions statement

H.T., H.C., I.T. and S.B. conceptualized the research. I.T. and H.C. designed the experiments. H.C. conducted the experiments, for which I.T. and H.C performed the analysis. S.B. supervised the research. All authors contributed to the writing, discussion, and revision of the manuscript.

## Acknowledgments

H.T. acknowledges funding support from the NSF graduate research fellowship program (GRFP) and Georgia Tech’s President’s Fellowship. S.B. acknowledges funding support from NIH Grant R35GM142588; NSF Grants MCB-1817334; CMMI-2218382; CAREER IOS-1941933; PHY-2310691 and the Open Philanthropy Project. Text in this paper was revised using LLMs.

## References

Baguñà, J., Romero, R., Saló, E., Collet, J., Auladell, C., and Bueno, D. (1990). Growth, degrowth and regeneration as developmental phenomena in adult freshwater planarians. Experimental embryology in aquatic plants and animals, pages 129–162.

Boddington, M. and Mettrick, D. (1974). The distribution, abundance, feeding habits, and population biology of the immigrant triclad dugesia polychroa (platyhelminthes: Turbellaria) in toronto harbour, canada. The Journal of Animal Ecology, pages 681–699.

Cliff, N. (1993). Dominance statistics: Ordinal analyses to answer ordinal questions. Psychological bulletin, 114(3):494.

Drewes, C. D. (1990). Tell-tail adaptations for respiration and rapid escape in a freshwater oligochaete (lumbriculus variegatus mull.). Journal of the Iowa Academy of Science: JIAS, 97(4):112–114.

Drewes, C. D. and Fourtner, C. R. (1989). Hindsight and rapid escape in a freshwater oligochaete. The Biological Bulletin, 177(3):363–371.

Hedrick, T. L. (2008). Software techniques for two- and three-dimensional kinematic measurements of biological and biomimetic systems. Bioinspiration & biomimetics, 3(3):034001.

Hyman, L. H. (1939). North american triclad turbellaria. ix. the priority of dugesia girard 1850 over euplanaria hesse 1897 with notes on american species of dugesia. Transactions of the American Microscopical Society, 58(3):264–275.

Kier, W. M. and Smith, K. K. (1985). Tongues, tentacles and trunks: the biomechanics of movement in muscular-hydrostats. Zoological journal of the Linnean Society, 83(4):307–324.

Kutschera, U. (2003). The feeding strategies of the leech Erpobdella octoculata (l.): A laboratory study. Int. Rev. Hydrobiol., 88(1):94–101.

Mann, H. B. and Whitney, D. R. (1947). On a test of whether one of two random variables is stochastically larger than the other. The annals of mathematical statistics, pages 50–60.

Miyamoto, M., Hattori, M., Hosoda, K., Sawamoto, M., Motoishi, M., Hayashi, T., Inoue, T., and Umesono, Y. (2020). The pharyngeal nervous system orchestrates feeding behavior in planarians. Science Advances, 6(15):eaaz0882.

Patil, V. P., Tuazon, H., Kaufman, E., Chakrabortty, T., Qin, D., Dunkel, J., and Bhamla, M. S. (2023). Ultrafast reversible self-assembly of living tangled matter. Science.

Pennak, R. (1978). Fresh-water invertebrates of the United States. Wiley, 2nd edition.

Perich, M. and Boobar, L. (1990). Effects of the predator dugesia dorotocephala (tricladida: Turbellaria) on selected nontarget aquatic organisms: Laboratory bioassay. Entomophaga, 35(1):79–83.

Pickavance, J. (1971). The diet of the immigrant planarian dugesia tigrina (girard): I. feeding in the laboratory. The Journal of Animal Ecology, pages 623–635.

Sluys, R. and Kawakatsu, M. (2005). Biodiversity of marine planarians revisited (platyhelminthes, tricladida, maricola). Journal of Natural History, 39(6):445–467.

Tuazon, H., David, S., Ma, K., and Bhamla, S. (2024). Leeches predate on fast-escaping and entangling blackworms by spiral entombment. Integrative and Comparative Biology, 64(5):1408–1415.

Tuazon, H., Kaufman, E., Goldman, D. I., and Bhamla, M. S. (2022). Oxygenation-controlled collective dynamics in aquatic worm blobs. Integrative and Comparative Biology, 62(4):890–896.

Tuazon, H., Nguyen, C., Kaufman, E., Tiwari, I., Bermudez, J., Chudasama, D., Peleg, O., and Bhamla, M. S. (2023). Collecting–Gathering biophysics of the blackworm lumbriculus variegatus. Integr. Comp. Biol., page icad080.

Vila-Farré, M. and C Rink, J. (2018). The ecology of freshwater planarians. Planarian regeneration: Methods and protocols, pages 173–205.

